# High-throughput detection and quantification of vitamin B_12_ in microbiome isolates using *Escherichia coli*

**DOI:** 10.1101/2025.04.08.647760

**Authors:** Katarzyna Hencel, Matthew Sullivan, Alper Akay

**Affiliations:** School of Biological Sciences, University of East Anglia, Norwich, NR4 7TJ, UK; Centre for Microbial Interactions, Norwich Research Park, Norwich, NR4 7UJ

**Keywords:** Vitamin B12, cobalamin, *Caenorhabditis elegans*, *C. elegans*, *E. coli*, *metE*, *CemBio*

## Abstract

Vitamin B_12_ is an essential micronutrient produced only by prokaryotes, and animals must acquire it from their diet. Vitamin B_12_ is critical for the synthesis of methionine and propionyl-CoA metabolism. In humans, vitamin B_12_ deficiency has been linked to many disorders, including infertility and developmental abnormalities. The growing trend towards plant-based diets and the ageing populations increase the risk of vitamin B_12_ deficiency, and therefore, there is an increasing interest in understanding vitamin B_12_ biology. Accurate approaches for detecting and quantifying vitamin B_12_ are essential in studying its complex biology, from its biogenesis in Bacteria and Archaea to its effects in complex organisms. Here, we present an approach using the commonly available *E. coli* methionine auxotroph strain B834 (DE3) and a multi-well spectrophotometer to detect and quantify vitamin B_12_ from biological samples at picomolar concentrations. We further show that our quantification method for vitamin B_12_ is sufficient to reveal important differences in the production of vitamin B_12_ from vitamin B_12_-synthesising bacteria commonly found in the microbiome of wild *Caenorhabditis elegans* isolates. Our results establish a high-throughput and simple assay platform for detecting and quantifying vitamin B_12_ using the *E. coli* B834 (DE3) strain.

## Introduction

Cobalamin (the natural form of vitamin B_12_) is, structurally, the most complex vitamin. It is only produced by bacteria and archaea and requires more than a dozen enzymes for its biogenesis. In many organisms, cobalamin (vitamin B_12_ hereafter) is essential for the function of two critical enzymes: methionine synthase, which regenerates methionine in the cells from homocysteine, and methylmalonyl-CoA mutase, which converts propionyl-CoA into succinyl-CoA. In humans, vitamin B_12_ deficiency has been linked to multiple diseases, including anaemia, infertility, and developmental and neurological disorders (1,2). Although clinical levels of vitamin B_12_ deficiency are rare (3), the global increase in plant-based diets and the ageing populations are linked to reduced vitamin B_12_ uptake, which is considered a growing global health risk that necessitates further molecular and medical research in vitamin B_12_ and its roles in human and animal physiology (4,5).

Research on vitamin B_12_ requires sensitive detection and quantification methods. Using vitamin B_12_ auxotrophy in bacteria to quantify vitamin B_12_ in biological samples has been a standard method since the 1940s when *Lactobacillus leichamnnii* was described by multiple groups as a suitable strain for vitamin B_12_ quantification (6–8). The assay has been used to this day and is readily available through commercial routes. However, *L. leichamnnii* has complicated growing conditions, including its response to thymidine in the absence of vitamin B_12_ (9).

Another bacterial strain commonly used for vitamin B_12_ assays is *Salmonella typhimurium* mutants, which lack the vitamin B_12_-independent methionine synthase, MetE (10). However, *Salmonella* strains grow under anaerobic conditions, often requiring additional apparatus. Another well-established assay uses the microalgae *Euglena gracilis var. bacillaris* (11). Although considered more accurate, this assay takes 4 to 7 days to complete and involves complicated growth conditions.

Alternatively, methionine auxotroph strains of *Escherichia coli (E. coli)* can also be used for vitamin B_12_ quantification (12–15). The advantages of utilising *E. coli* strains for vitamin B_12_ assays include fast growth, which allows the assay to be performed overnight, and simple growth requirements, which do not involve specialised media preparation. However, there is limited information on the sensitivity and specificity of *E. coli*-based vitamin B_12_ assays.

Here, we present a high-throughput bacteria-based assay for vitamin B_12_ detection and quantification using the methionine-auxotroph *E. coli* B834 (DE3) strain. This approach allows for simple and time-efficient detection and quantification of vitamin B_12_ content in biological samples. We further utilise the method to quantify the vitamin B_12_ content of different bacterial species commonly found with wild isolates of the nematode *Caenorhabditis elegans (C. elegans)*.

## Methods

### Bacterial strains, plasmids and culture media

All bacteria and plasmids are listed in Table 2 and Table 3, respectively. *E. coli* OP50, *C. aquatica* DA1877 and the isogenic Δ*cbiA*Δ*cbiB* mutant were grown in the soya-rich medium at 37 °C at 180 rpm agitation for the extract preparation. *E. coli* B834 (DE3) was grown in M9 minimal salts medium (KH_2_PO_4_, 15 g/L NaCl, 2.5 g/L Na_2_HPO_4_, 33.9 g/L NH_4_Cl, 5 g/L, 2 mM MgSO_4_, 0.1 mM CaCl_2_, 0.4 % glucose unless otherwise stated) supplemented with 400 nM L-methionine at 37 °C at 180 rpm. Strains from the CeMbio collection for the extract preparation were grown at 28 °C at 180 rpm in the vitamin B12-deficient, soya-rich medium (soya peptone 20 g/L, sodium chloride 5 g/L).

**Table 1.**
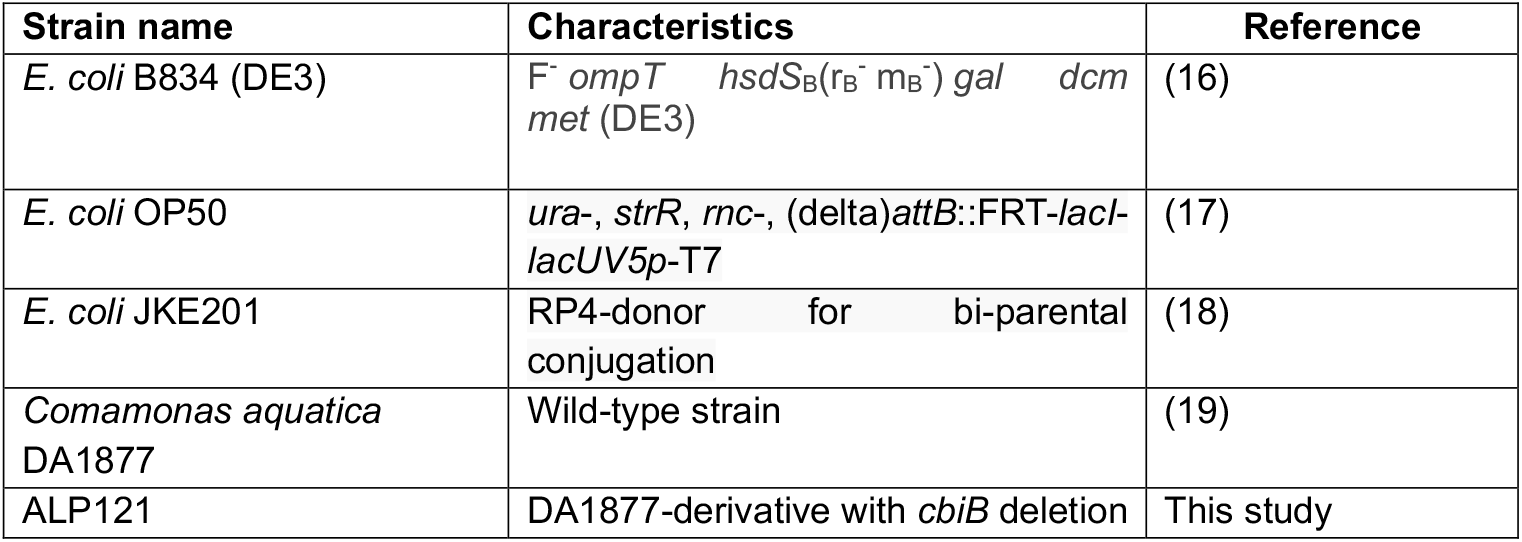
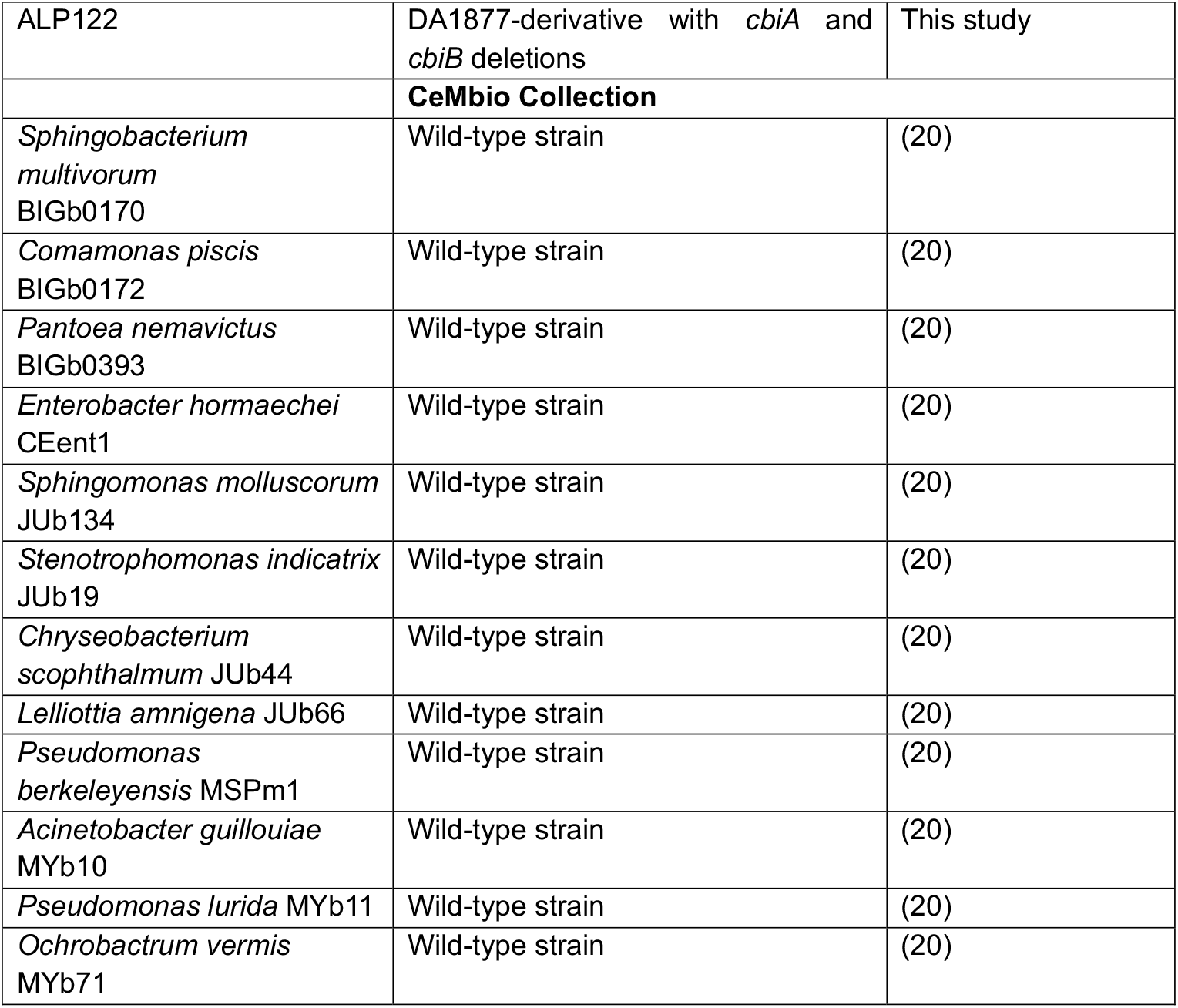
Bacterial strains used in the study.

**Table 2.** Plasmids used in this study.

| Plasmid | Description | Reference |
| --- | --- | --- |
| pFOK | Suicide vector containing *sacB* and driven by *ptetA*; kan<sup>R</sup> | (21) |
| pAA46 | Derivative of pFOK containing up- and downstream fragments of the *cbiB*; kan<sup>R</sup> | This study |
| pAA47 | Derivative of pFOK containing up- and downstream fragments of the *cbiA*; kan<sup>R</sup> | This study |

**Table 3.**
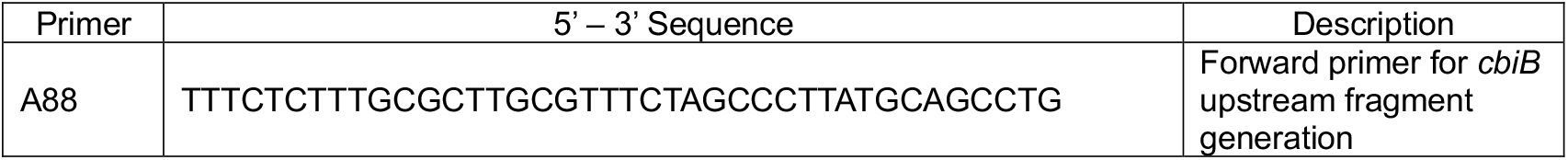
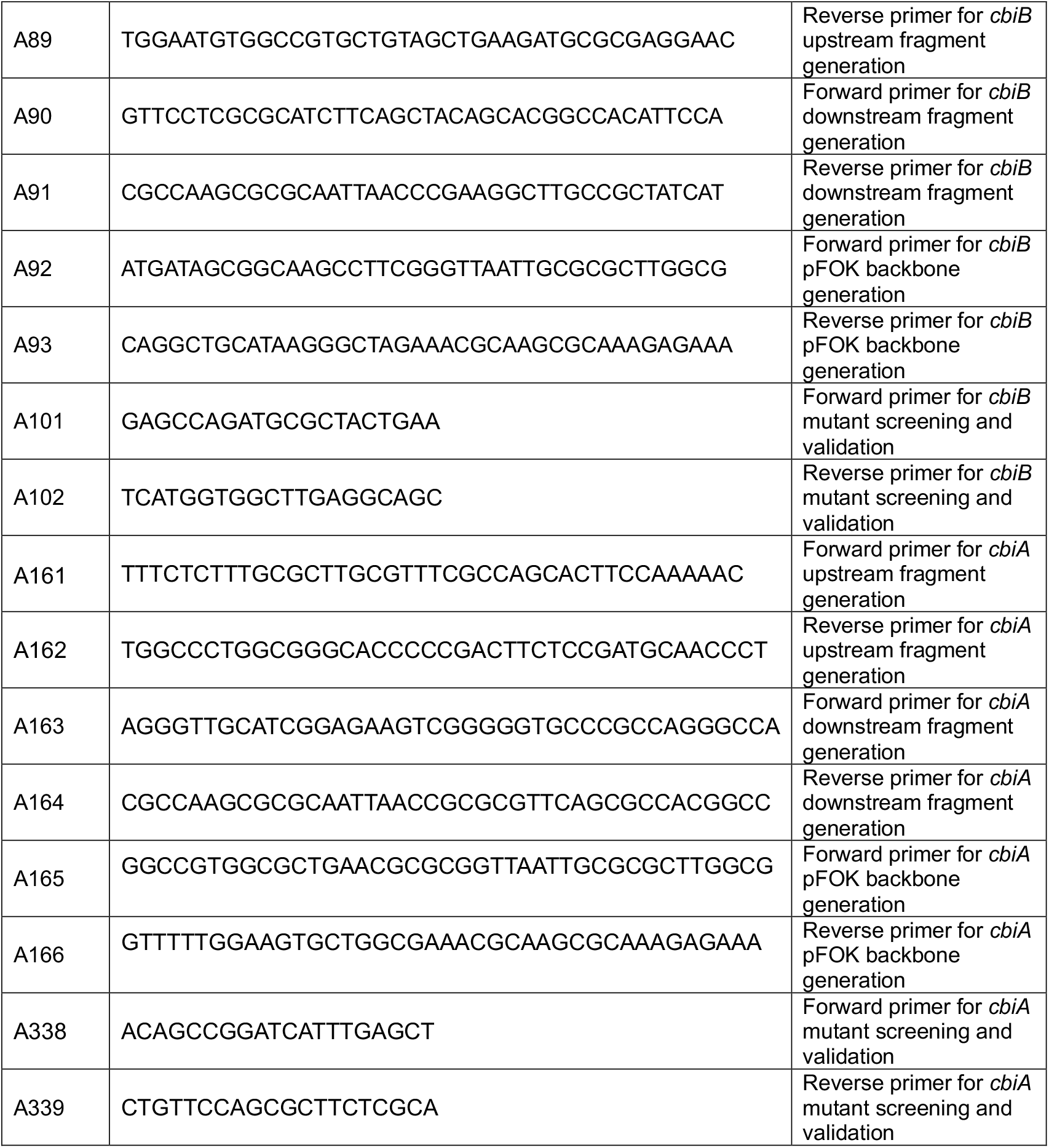
Primers used in the study.

### *C. aquatica* DA1877 vitamin B_12_-deficient mutant generation

Oligonucleotide primers (Table 3) with flanking 20 bp overhangs were designed to amplify upstream and downstream fragments from *cbiA* and *cbiB* of *C. aquatica* DA1877 using Benchling’s Gibson Assembly Wizard. The amplified fragments were introduced to the pFOK suicide vector through Gibson Assembly and transformed into *E. coli* JKE201. All constructs were verified by Sanger sequencing, followed by conjugation with *C. aquatica* DA1877 on LB supplemented with 100 µM diaminopimelic acid (DAP) to support *E. coli* JKE201 growth. Transconjugants were selected onto LB agar containing 100 µg/mL kanamycin. At least 3 transconjugants were grown in LB medium for 4 hours, followed by plating on no-salt LB agar plates (10 g/L tryptone, 5 g/L yeast extract, 15 g/L agar) supplemented with 20 % sucrose and 0.5 µg/mL anhydro-tetracycline. Candidate colonies were screened for deletions through PCR and verified by Sanger sequencing. Mutation in *cbiB* resulted in the out-of-frame deletion of 178 amino acids, while *cbiA* mutation resulted in complete gene removal. The Sanger sequencing trace files are available as Supplementary File 1.

### Bacterial lysate preparation

Bacterial cultures for the *E. coli* B834 (DE3) assay were grown overnight and 1 OD unit (equivalent of 1 mL of culture with absorbance at 600nm of 1.0) was centrifuged at 15,000 rpm for 1 minute. The supernatant was removed, and the cells were resuspended in 50 µL of M9 minimal salts medium and boiled at 100 °C for 15 minutes, as previously described (22). After boiling, lysates were centrifuged at 15,000 rpm for 1 minute to remove debris, and the cooled supernatant was used as an extract for supplementation assays.

### *E. coli* B834 (DE3) Assay

The assay was prepared in 96-well plates (*Greiner* #655180) with the final volume of 200 µL of M9 minimal salts medium. Overnight cultures of *E. coli* B834 (DE3) grown in LB were back-diluted 1:100 into the wells and supplemented with either 2 µL of prepared bacterial lysates (extracts) or vitamin B_12_ standard solutions used for the growth curves. The growth response was recorded over 20 hours at 37°C with 300 rpm agitation, with readings taken every 30 minutes using a SPECTROstar*®* Nano plate reader (BMG Labtech) in matrix scan mode using a 2x2 scan matrix with 25 flashes per scan point and path length correction of 5.88mm for 200 µL volume. For blank corrections of optical density readings, control wells containing media without bacteria were included. Methylcobalamin (Thermo Scientific Chemicals, #A11176ME) was used for the vitamin B_12_ standard curve.

### Visualisation and analysis

Data was analysed and visualised using Prism 10 (Version 10.3.0).

## Results

### Analysis of *E. coli* B834 methionine and vitamin B_12_ auxotrophy

*E. coli* has two methionine synthase enzymes: the B_12_-dependent MetH and the B_12_-independent MetE. *E. coli* B834 (DE3) has a *null* mutation in the *metE* gene, making the bacteria solely dependent on either methionine or B_12_ supplementation. To assess whether *E. coli* B834 (DE3) is suitable for vitamin B_12_ detection and quantification, we tested the growth of this strain in response to various vitamin B_12_ and methionine supplementations. We confirmed that *E. coli* B834 (DE3) can grow only when methionine or vitamin B_12_ is present in the media (Figure 1A). We did not observe any significant difference in growth when B_12_ was supplemented at concentrations ranging from 1 nM to 1000 nM, as determined by area under the curve analysis followed by one-way ANOVA with Holm-Sidak multiple comparison corrections (Figure 1B).

**Figure 1.**
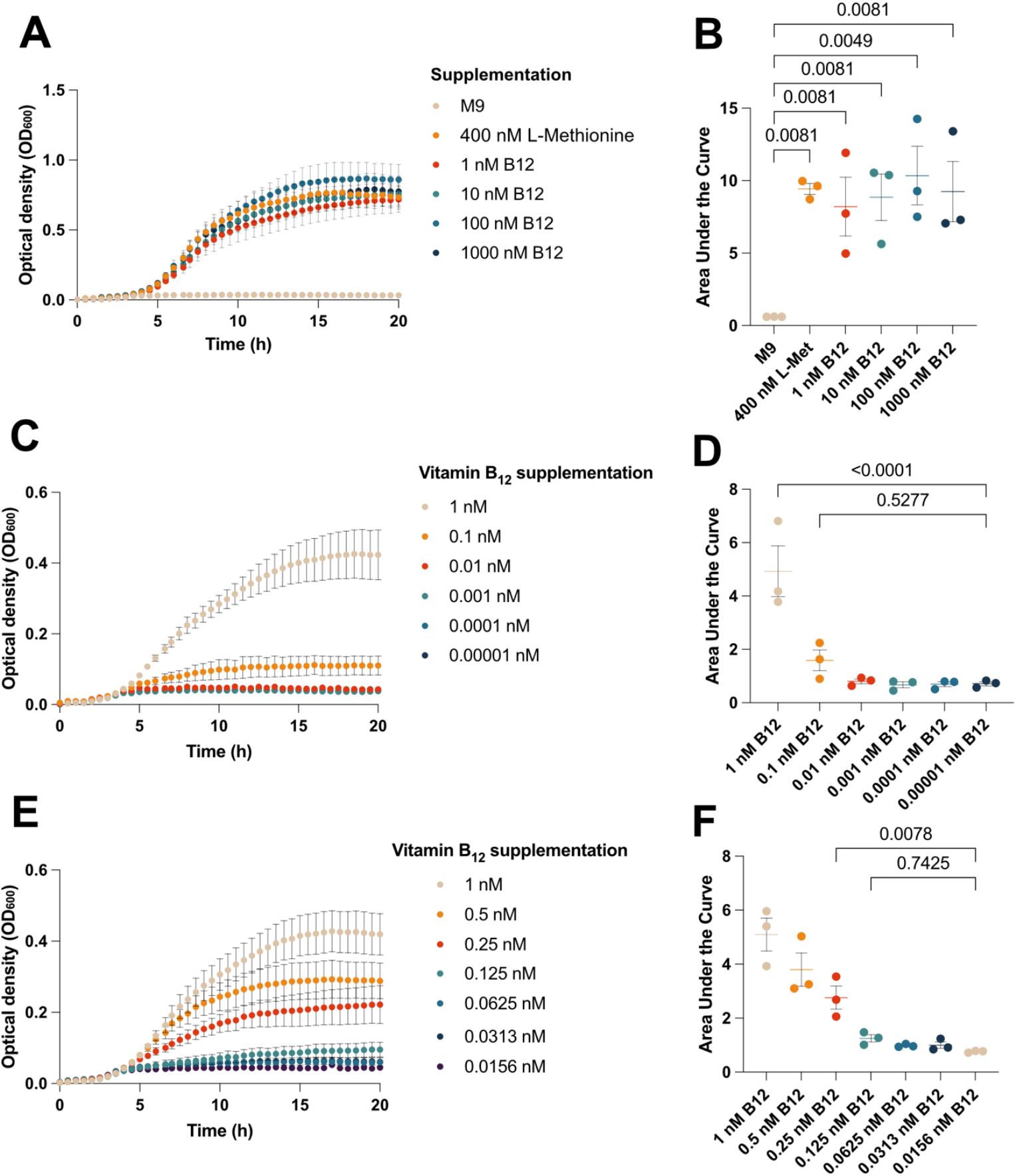
The growth of *E. coli* B834 is dependent on methionine or vitamin B_12_ and is titratable with vitamin B_12_ concentration. *E. coli* B834 was cultured in M9 minimal media, without or with supplemental methionine or vitamin B_12_ (1 nM – 1000 nM) as indicated **(A)**. The growth conditions were compared by Area Under the Curve analysis, followed by one-way ANOVA with Holm-Šidák multiple comparison corrections and P-values indicated **(B)**. *E. coli* B834 growth with vitamin B_12_ supplementation at 1 nM and using 10-fold **(C)** and 2-fold serial dilutions **(E)** to test a broad range of sub-nanomolar vitamin B_12_ concentrations. The growth of *E. coli* B834 in the 10-fold and 2-fold were compared by Area Under the Curve analysis followed by one-way ANOVA and Holm-Šidák multiple comparison corrections and comparisons are shown **(D)** and **(F)**, respectively. The data points are plotted as mean ± SEM from 3 biological replicates derived from 2 technical replicates.

### Using the growth of *E. coli* B834 as a measure of vitamin B_12_ quantity

We subsequently sought to test the utility of using the growth of *E. coli* B834 (DE3) as a highly sensitive biological method for detecting vitamin B_12_. To this end, we used M9 minimal media (devoid of methionine or vitamin B_12_) and added vitamin B_12,_ supplemented at concentrations ranging from 0.00001 nM to 1 nM using 10-fold (Figures 1C and 1D) and 2-fold (Figures 1E and 1F) serial dilutions. Using this range, we determined that vitamin B_12_ concentrations at and above 0.25 nM (250 pM) were sufficient to support the growth of the *E. coli* B834 (DE3) strain, as evidenced by a significant increase in area under the curve measurements throughout the growth period (Figure 1F). Next, we prepared a standard curve of vitamin B_12_ concentrations between 0 and 0.4 nM to determine the limit of detection and quantification for vitamin B_12_ using the *E. coli* B834 (DE3) strain. Compared to the unsupplemented media control, the limit of detection is 0.05 nM (50 pM) (Figure 2A and 2B), indicating that the growth of *E. coli* was detectable above background absorbance measurements. Increasing the carbon source from 0.4% glucose to 1.0% did not change the sensitivity of the growth assay (Supplementary Figure 1A and 1B). In summary, we have established that the growth of *E. coli* B834 (DE3) can be used to detect and quantify vitamin B_12_ at picomolar concentrations.

**Figure 2.**
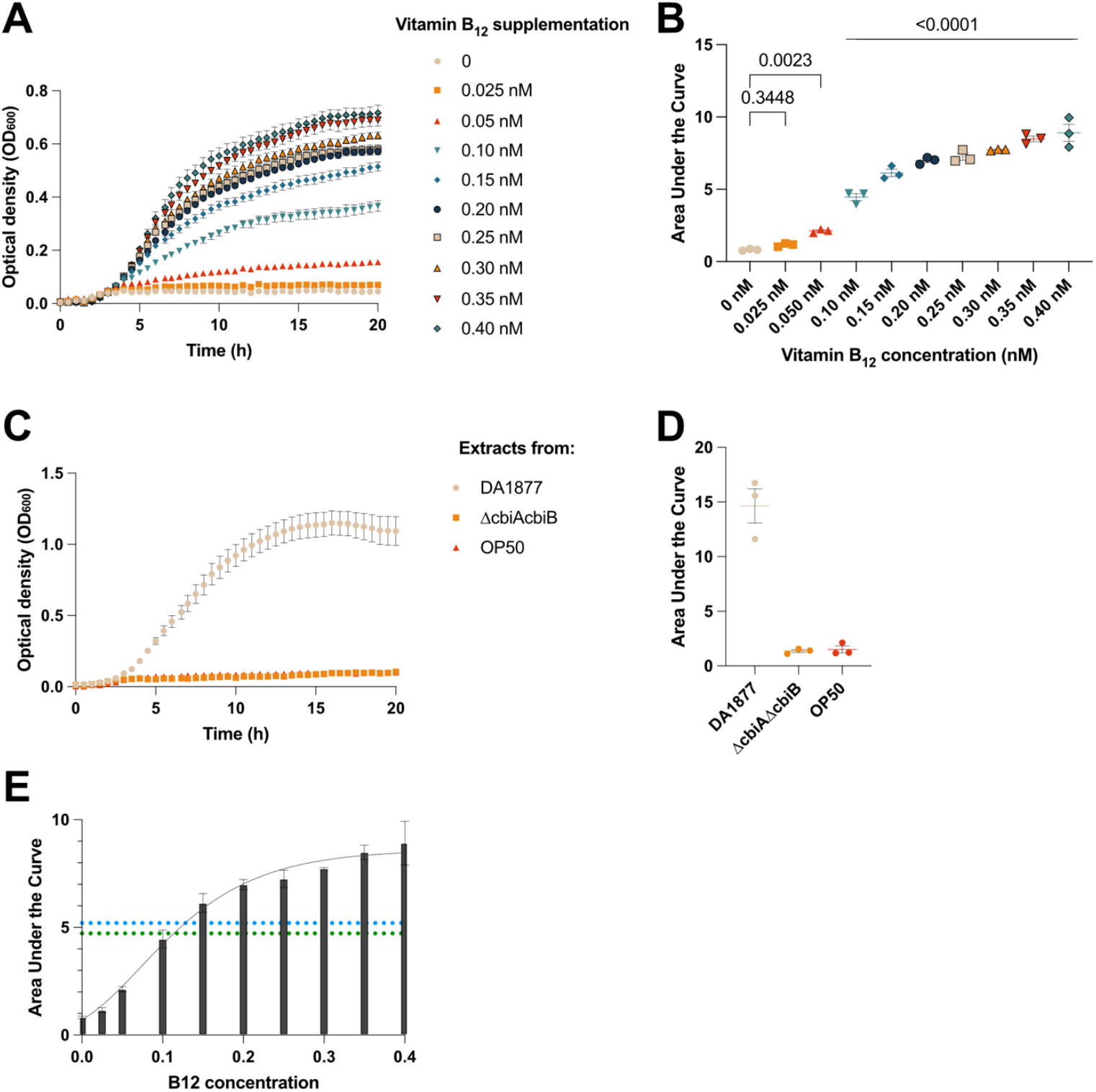
Use of *E. coli* B834 growth to detect and quantify vitamin B_12_ production by *C. aquatica*. *E. coli* B834 was cultured in M9 minimal medium, without or with supplemental vitamin B_12_ at concentrations between 0.025 nM – 0.4 nM (**A**) and compared using Area under the Curve analysis coupled with one-way ANOVA with Holm-Šidák multiple comparison corrections **(B)**. *E. coli* B834 culture was supplemented by bacterial cell-free extracts of *C. aquatica* DA1877, Δ*cbiA*Δ*cbiB C. aquatica*, and *E. coli* OP50 **(C)** and growth in the presence of extracts were compared by Area under the Curve analysis coupled with one-way ANOVA with Holm-Šidák multiple comparison corrections **(D)**. Area under the Curve data from vitamin B_12_ standards in Panel B were used for Gompertz model fitting, further employed for vitamin B_12_ quantification (*Y = YM*(Y0/YM)^(exp(-K*X)*), where YM is the maximum AUC score, Y0 is the minimum AUC score, K determines the lag time. The dashed green line corresponds to the 4.70 AUC score of the 10^-1^ dilution of *C. aquatica* DA1877 extract, while the dashed blue line corresponds to the 5.20 AUC score of the 1:8 dilution of *C. aquatica* DA1877 extract, where 2 µL out of 50 µL of the 1 OD extract was used for quantification. In this model, YM = 8.606, Y0 = 0.7062, and K = 12.53. All data points are plotted with mean ± SEM from 3 biological replicates derived from 2 technical replicates.

### Using *E. coli* B834 (DE3) to quantify vitamin B_12_ in biological samples

*C. elegans* is a well-established model organism for studying the function of vitamin B_12_ during animal development and for understanding the molecular pathways related to vitamin B_12_ (24–28). *C. elegans* exclusively feeds on bacteria, and its uptake of vitamin B12 depends on the bacteria available in its environment as a food source. One such bacterium *C. elegans* feeds on in the wild is *Comamonas aquatica (C. aquatica)* DA1877, a known vitamin B_12_ producer (28) that was isolated from soils (19). Mutations in the *cbiA* and *cbiB* genes, which code for cobyrinate a,c-diamide synthase and adenosylcobinamide-phosphate synthase enzymes, respectively, prevent *C. aquatica* from producing vitamin B_12_ (28). As a negative control for our assay, we generated an isogenic Δ*cbiA*Δ*cbiB* mutant of DA1877 by deleting the *cbiA* and *cbiB* genes (Supplementary Figure 2A-2C). To confirm that our Δ*cbiA*Δ*cbiB* strain no longer produced vitamin B_12_, we utilised our *E. coli* B834 (DE3) approach to test for the presence of vitamin B_12_ in cell-free extracts from *C. aquatica*. Briefly, bacterial cells were lysed by boiling, and cell-free extracts were added to *E. coli* B834 (DE3) in media devoid of vitamin B_12_ or methionine. Using this approach, we assayed the vitamin B_12_ levels of wild-type *C. aquatica, C. aquatica* Δ*cbiA*Δ*cbiB* and *E. coli* OP50, a different strain of *E. coli* commonly used as laboratory food for *C. elegans* but known to be a vitamin B_12_ non-producer (28,29). As predicted, cell-free extracts of wild-type *C. aquatica* DA1877 supported vitamin B_12_-dependent growth of *E. coli* B834 (DE3), whereas the *C. aquatica* Δ*cbiA*Δ*cbiB* mutant and *E. coli* OP50 did not (Figure 2C and 2D). We further quantified the vitamin B_12_ content from *C. aquatica* extracts by using 2-fold and 10-fold serial dilutions and comparing the relative growth of *E. coli* B834 (DE3) with *C. aquatica* extracts against growth with known concentrations of vitamin B_12_ (Supplementary Figure 2D and 2E). Using the 2-fold and 10-fold dilutions combined with the Gompertz-modelled Area under the Curve analysis of the standard curve, we estimate the vitamin B_12_ content of *C. aquatica* DA1877 to be approximately 25 nM per 1 OD unit (1 mL of culture at 1.0 OD_600nm_) of bacteria (Figure 2E).

Our assays with vitamin B12, along with extracts from both vitamin B12-producing and non-producing bacteria, provided proof of concept for our method to detect vitamin B12 in complex biological samples by using the growth of E. coli B834 (DE3) as a proxy. Next, we applied this method to assess the vitamin B12 content of 12 bacterial strains from the CeMbio collection, all of which were isolated from *C. elegans* found in the wild (20). Four bacterial strains, *Comamonas piscis* BIGb0172, *Pseudomonas berkeleyensis* MSPm1, *Pseudomonas lurida* MYb11 and *Ochrobactrum vermis* MYb71, were predicted to produce vitamin B_12_ based on their genomic sequences and predicted metabolic pathway analyses (20,29). Our analysis using the *E. coli* B834 (DE3) growth assay showed that *C. piscis* BIGb0172, *P. berkeleyensis* MSPm1, *P. lurida* MYb11, and *O. vermis* MYb71 are indeed vitamin B12 producers, because cell-free extracts from cultures of these bacteria were capable of supporting the growth of *E. coli* B834 (DE3) in a manner that relied on vitamin B12 (Figure 3A and 3B). Among these, extracts from *C. piscis* BIGb0172 and *P. berkeleyensis* MSPm1 supported the highest growth, while *O. vermis* MYb71 showed reduced growth, indicating that there may be variation in the amount of vitamin B_12_ produced by these bacteria. In contrast, supplementing *E. coli* B834 (DE3) with extracts from the other eight CeMbio strains led to a complete absence of bacterial growth (Figure 3A and 3B).

**Figure 3.**
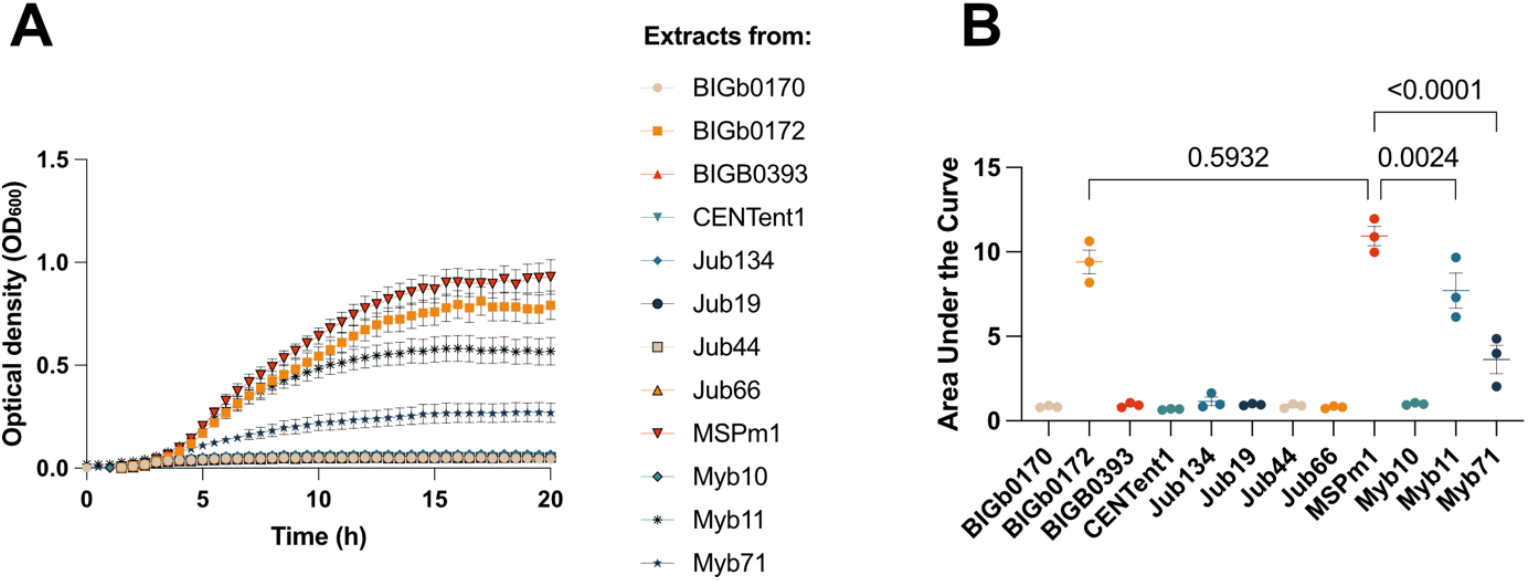
Quantification of vitamin B_12_ levels in bacterial isolates from wild *C. elegans*. *E. coli* B834 was cultured in M9 minimal media and supplemented with cell-free bacterial extracts of the indicated strains **(A)**. Growth was compared using Area Under the Curve analysis coupled with one-way ANOVA with Holm-Šidák multiple comparison corrections **(B)**. The data points are plotted with mean ± SEM from 3 biological replicates derived from 2 technical replicates corrected for blank readings. Bacterial isolates are as follows: *Sphingobacterium multivorum* BIGb0170, *Comamonas piscis* BIGb0172, *Pantoea nemavictus* BIGb0393, *Enterobacter hormaechei* CEent1, *Sphingomonas molluscorum* JUb134, *Stenotrophomonas indicatrix* JUb19, *Chryseobacterium scophthalmum* JUb44, *Lelliottia amnigena* JUb66, *Pseudomonas berkeleyensis* MSPm1, *Acinetobacter guillouiae* MYb10 *Pseudomonas lurida* MYb11, *Ochrobactrum vermis* MYb71.

In summary, we show that our application of *E. coli* B834 (DE3) growth in minimal media can be used for rapid and high-throughput detection and relative quantification of vitamin B_12_ in biological samples.

## Discussion

Vitamin B_12_-dependent microorganisms are commonly used to detect and quantify vitamin B_12_ in various formats (6–8). Using *E. coli metE* mutants for this purpose offers numerous advantages, including their commercial availability, rapid growth, and simple growth requirements. However, there is limited information on the sensitivity and reproducibility of *E. coli metE*-based vitamin B_12_ assays. Here, we present a vitamin B_12_ quantification assay using a readily available commercial strain of *E. coli* B834 (DE3) and widely used and inexpensive minimal media. The assay was developed in liquid culture using a 96-well plate format and a multi-well plate reader, allowing for reproducible analysis of many biological samples with a sensitivity as low as 50 picomolar concentration.

The dependency on the growth of *E. coli* B834 (DE3) due to the presence of vitamin B_12_ can be confounded by the presence of methionine, which may bypass the metabolic bottleneck caused by vitamin B_12_ limitation in *metE*^-^ *E. coli* and could affect the specificity of our assay. However, previous studies conducted on the *E. coli* 113-3 strain, another methionine auxotroph, showed that methionine must be 50,000 times more concentrated than vitamin B_12_ to hinder vitamin B_12_ quantification using the *E. coli* assay, which was significantly higher than the levels found in the mammalian tissues tested (12). Similarly, we did not observe unexpected *E. coli* B834 (DE3) growth supported by extracts derived from known vitamin B_12_ non-producers (Figure 2C) or CeMbio collection strains, which were not predicted to produce vitamin B_12_ (Figure 3).

Previous studies suggested that four strains in the CeMbio collection, *C. piscis* BIGb0172, *P. berkeleyensis* MSPm1, *P. lurida*, and *O. vermis* Myb71, are vitamin B_12_ producers, based on the presence of vitamin B_12_ biosynthetic pathway genes (29). Our analysis provided experimental evidence to support this. Interestingly, despite confirming that all four isolates are vitamin B_12_ producers, we note that the levels of vitamin B_12_ likely vary significantly, with *P. berkeleyensis* MSPm1 and *C. piscis* BIGb0172 producing significantly higher levels of vitamin B_12_ compared to *P. lurida* Myb11 and *O. vermis* Myb71. These differences in vitamin B_12_ content could be important for the growth of *C. elegans* and other organisms that directly rely on bacteria for vitamin B_12_.

## Conclusions

Our results establish a high-throughput, straightforward, and cost-effective method for detecting and quantifying vitamin B_12_ levels in biological samples. The simplicity, reproducibility, and sensitivity of the *E. coli* B834 (DE3) assay provide an important methodology for the research community working on vitamin B_12_. Our discovery of varying vitamin B_12_ levels in the wild *C. elegans* microbiome makes a compelling case for further investigation into how differences in bacterial metabolites impact animal development.

## Ethics approval and consent to participate

Not applicable.

## Consent for publication

Not applicable.

## Availability of data and materials

All data generated or analysed during this study are included in this published article [and its supplementary information files].

## Author’s contributions

Katarzyna Hencel, Matthew Sullivan and Alper Akay contributed to the conceptualisation, data analysis and manuscript writing. Katarzyna Hencel carried out the data acquisition and wrote the first draft of the manuscript. Alper Akay acquired the funding for the work.

## Acknowledgements

We would like to thank UEA School of Biological Sciences technicians and the admin team for their support throughout the project. We thank the Caenorhabditis Genetics Centre (CGC), funded by the NIH Office of Research Infrastructure Programs (P40 OD010440). We thank Wormbase for providing access to essential *C. elegans* resources.

## Supplementary Figure Legends

**Supplementary Figure 1.**
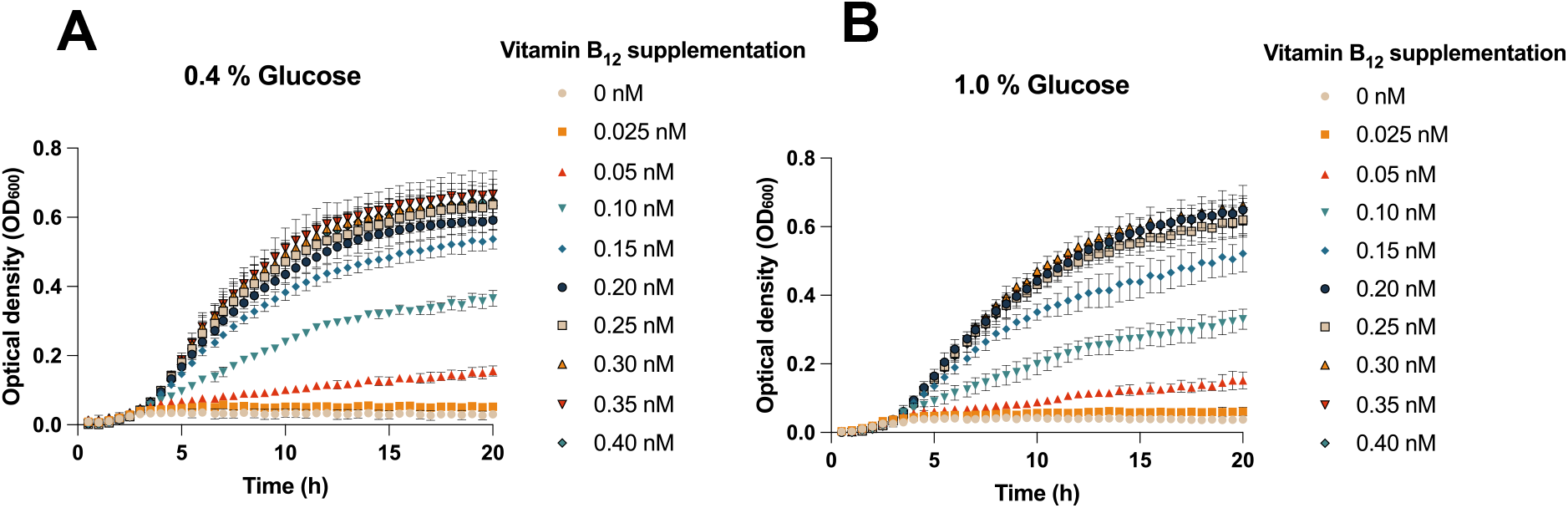
*E. coli* B834 was cultured in M9 minimal media with vitamin B_12_ supplementations of 0.025 nM – 0.40 nM under standard carbon source input (0.4 % glucose; **A**) or increased carbon source input (1.0 % glucose; **B**). The data points are plotted with mean ± SEM from 3 biological replicates derived from 2 technical replicates corrected for blank readings.

**Supplementary Figure 2.**
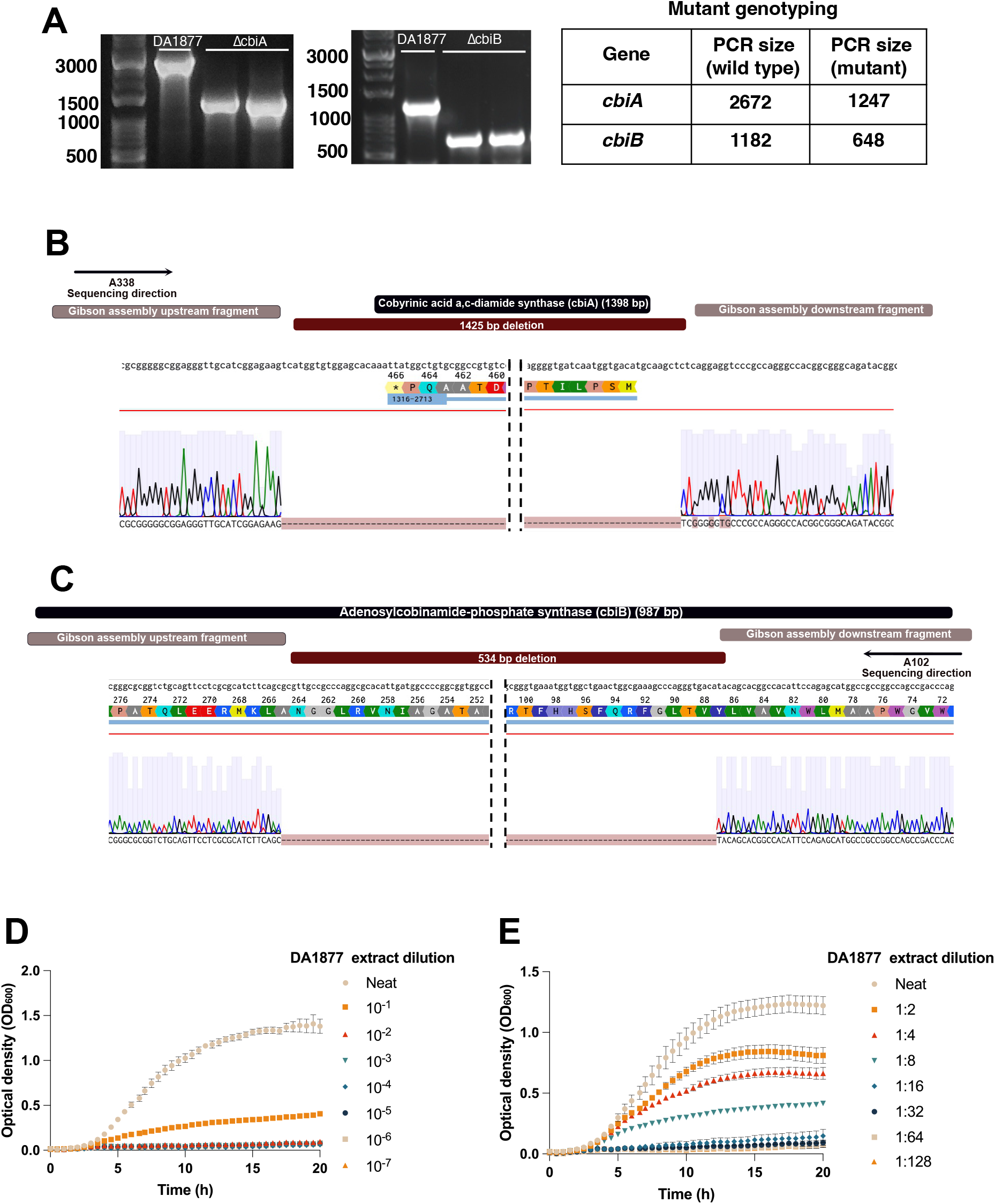
**(A)** PCR validation of *cbiA* and *cbiB* deletions. For the *cbiA* gene deletion validation, primers A338 and A339 were used, with the wild type *C. aquatica* DA1877 genotype of 2672 bp and the length of 1274 bp of a successful *cbiA* mutant. For the *cbiB* mutation validation, primers A101 and A102 were used, with the wild type *C. aquatica* DA1877 genotype of 1182 bp and the length of 648 bp of a successful *cbiB* mutant. PCR validation was followed by the **(B-C)** Sanger sequencing validation using A338 and A102 primers for *cbiA* and *cbiB* genes, respectively, aligned against the reference genome of *C. aquatica* DA1877. The red dashed region of the sequencing profiles corresponds to the deleted portions of both genes and the lack of sequencing signal. The vertical dashed lines represent the cropped region of the deleted portions of the genes. **(D - E)** Vitamin B_12_ quantification in *C. aquatica* DA1877 was performed by supplementing *C. aquatica* DA1877 extracts in **(D)** 10-fold and **(E)** 2-fold dilutions to *E. coli* B834. The data points are plotted with mean ± SEM from 3 biological replicates derived from 2 technical replicates corrected for blank readings.

## References

1. Mischoulon D, Burger JK, Spillmann MK, Worthington JJ, Fava M, Alpert JE. Anemia and macrocytosis in the prediction of serum folate and vitamin B12 status, and treatment outcome in major depression. Journal of Psychosomatic Research. 2000 Sep 1;49(3):183– 7.

2. Molloy AM, Kirke PN, Brody LC, Scott JM, Mills JL. Effects of Folate and Vitamin B12 Deficiencies During Pregnancy on Fetal, Infant, and Child Development. Food Nutr Bull. 2008 Jun 1;29(2_suppl1):S101–11.

3. Green R, Allen LH, Bjørke-Monsen AL, Brito A, Guéant JL, Miller JW, et al. Vitamin B12 deficiency. Nat Rev Dis Primers. 2017 Jun 29;3(1):17040.

4. Brouwer-Brolsma EM, Dhonukshe-Rutten RAM, Van Wijngaarden JP, Zwaluw NL van der, Velde NV der, De Groot LCPGM. Dietary Sources of Vitamin B-12 and Their Association with Vitamin B-12 Status Markers in Healthy Older Adults in the B-PROOF Study. Nutrients. 2015 Sep;7(9):7781–97.

5. Niklewicz A, Smith AD, Smith A, Holzer A, Klein A, McCaddon A, et al. The importance of vitamin B12 for individuals choosing plant-based diets. Eur J Nutr. 2023 Apr;62(3):1551–9.

6. Hoffmann CE, Stokstad ELR, Hutchings BL, Dornbush AC, Jukes TH. THE MICROBIOLOGICAL ASSAY OF VITAMIN B12 WITH LACTOBACILLUS LEICHMANNII. Journal of Biological Chemistry. 1949 Dec 1;181(2):635–44.

7. Hoffmann CE, Stokstad ELR. Response of Lactobacillus leichmannii 313 to the antipernicious anemia factor. J Biol Chem. 1948 Dec;176(3):1465.

8. Skeggs HR, Huff JW. The use of Lactobacillus leichmannii in the microbiological assay of the animal protein factor. J Biol Chem. 1948 Dec;176(3):1459.

9. Kitay E, McNutt WS, Snell EE. Desoxyribosides and vitamin b12 as growth factors for lactic acid bacteria. Journal of Bacteriology. 1950 Jun;59(6):727–38.

10. Raux E, Lanois A, Levillayer F, Warren MJ, Brody E, Rambach A, et al. Salmonella typhimurium cobalamin (vitamin B12) biosynthetic genes: functional studies in S. typhimurium and Escherichia coli. J Bacteriol. 1996 Feb;178(3):753–67.

11. Hutner SH, Provasoli L. Assay of anti-pernicious anemia factor with Euglena. Proc Soc Exp Biol Med. 1949 Jan;70(1):118–20.

12. Chiao JS, Peterson WH. Microbiological Assay of Vitamin B12 with a Mutant Strain of Escherichia coli. Appl Microbiol. 1953 Jan;1(1):42–6.

13. Davis BD, Mingioli ES. Mutants of Escherichia coli requiring methionine or vitamin B12. J Bacteriol. 1950 Jul;60(1):17–28.

14. Harrison E, Lees KA, Wood F. The assay of vitamin B12. Part VI. Microbiological estimation with a mutant of Escherichia coli by the plate method. Analyst. 1951 Jan 1;76(909):696–705.

15. Johansson KR. Response to and Assay of Vitamin B12 by a Mutant of Escherichia coli. Proceedings of the Society for Experimental Biology and Medicine. 1953 Jul 1;83(3):448– 53.

16. Wood WB. Host specificity of DNA produced by Escherichia coli: bacterial mutations affecting the restriction and modification of DNA. J Mol Biol. 1966 Mar;16(1):118–33.

17. Brenner S. The Genetics of CAENORHABDITIS ELEGANS. Genetics. 1974 May;77(1):71–94.

18. Harms A, Liesch M, Körner J, Québatte M, Engel P, Dehio C. A bacterial toxin-antitoxin module is the origin of inter-bacterial and inter-kingdom effectors of Bartonella. PLoS Genet [Internet]. 2017 Oct 26 [cited 2021 May 1];13(10). Available from: https://www.ncbi.nlm.nih.gov/pmc/articles/PMC5675462/

19. Shtonda BB, Avery L. Dietary choice behavior in Caenorhabditis elegans. J Exp Biol. 2006 Jan;209(Pt 1):89–102.

20. Dirksen P, Assié A, Zimmermann J, Zhang F, Tietje AM, Marsh SA, et al. CeMbio - The Caenorhabditis elegans Microbiome Resource. G3 (Bethesda). 2020 Sep 2;10(9):3025–39.

21. Cianfanelli FR, Cunrath O, Bumann D. Efficient dual-negative selection for bacterial genome editing. BMC Microbiol. 2020 May 24;20:129.

22. Ross GIM. Vitamin B12 Assay in Body Fluids using Euglena gracilis. J Clin Pathol. 1952 Aug;5(3):250–6.

23. Frederick RO, Bergeman L, Blommel PG, Bailey LJ, McCoy JG, Song J, et al. Small-scale, semi-automated purification of eukaryotic proteins for structure determination. J Struct Funct Genomics. 2007 Dec 1;8(4):153–66.

24. Bito T, Okamoto N, Otsuka K, Yabuta Y, Arima J, Kawano T, et al. Involvement of Spermidine in the Reduced Lifespan of Caenorhabditis elegans During Vitamin B12 Deficiency. Metabolites. 2019 Sep;9(9):192.

25. Bito T, Misaki T, Yabuta Y, Ishikawa T, Kawano T, Watanabe F. Vitamin B12 deficiency results in severe oxidative stress, leading to memory retention impairment in Caenorhabditis elegans. Redox Biol. 2017 Apr;11:21–9.

26. Bito T, Matsunaga Y, Yabuta Y, Kawano T, Watanabe F. Vitamin B12 deficiency in Caenorhabditis elegans results in loss of fertility, extended life cycle, and reduced lifespan. FEBS Open Bio. 2013;3:112–7.

27. Bito T, Watanabe F. Biochemistry, function, and deficiency of vitamin B12 in Caenorhabditis elegans. Exp Biol Med (Maywood). 2016 Sep;241(15):1663–8.

28. Watson E, MacNeil LT, Ritter AD, Yilmaz LS, Rosebrock AP, Caudy AA, et al. Interspecies systems biology uncovers metabolites affecting C. elegans gene expression and life history traits. Cell. 2014 Feb 13;156(4):759–70.

29. Zimmermann J, Obeng N, Yang W, Pees B, Petersen C, Waschina S, et al. The functional repertoire contained within the native microbiota of the model nematode Caenorhabditis elegans. ISME J. 2020 Jan;14(1):26–38.

